# 5-Hydroxymethylcytosines from Circulating Cell-free DNA as Diagnostic and Prognostic Markers for Hepatocellular Carcinoma

**DOI:** 10.1101/424978

**Authors:** Jiabin Cai, Lei Chen, Zhou Zhang, Xinyu Zhang, Xingyu Lu, Weiwei Liu, Guoming Shi, Yang Ge, Pingting Gao, Yuan Yang, Aiwu Ke, Linlin Xiao, Ruizhao Dong, Yanjing Zhu, Xuan Yang, Jiefei Wang, Tongyu Zhu, Deping Yang, Xiaowu Huang, Chengjun Sui, Shuangjian Qiu, Feng Shen, Huichuan Sun, Weiping Zhou, Jian Zhou, Ji Nie, Xu Zhang, Brian C-H.Chiu, Wan Yee Lau, Chuan He, Hongyang Wang, Wei Zhang, Jia Fan

## Abstract

The lack of highly sensitive and specific diagnostic biomarkers is a major contributor to the poor outcomes of patients with hepatocellular carcinoma (HCC), the second-most common cause of cancer deaths worldwide. We sought to develop a clinically convenient and minimally-invasive approach that can be deployed at scale for the sensitive, specific, and highly reliable diagnosis of HCC, and to evaluate the potential prognostic value of this approach. The study cohort comprised of 2,728 subjects, including HCC patients (n = 1,208), controls (n = 965) (572 healthy individuals and 393 patients with benign lesions), as well as patients with chronic hepatitis B infection (CHB) (n =291), liver cirrhosis (LC) (n = 110), and cholangiocarcinoma (CCC) (n = 154), was recruited from three major liver cancer hospitals in Shanghai, China, from July 2016 to November 2017. Circulating cell-free DNA (cfDNA) were collected from plasma samples from these individuals before surgery or any radical treatment. Applying our 5hmC-Seal technique, the summarized 5-hydroxymethylcytosine (5hmC) profiles in cfDNA were obtained. Molecular annotation analysis suggested that the profiled 5hmC loci in cfDNA were enriched with liver tissue-derived regulatory markers (e.g., H3K4me1). We showed that a weighted diagnostic score (wd-score) based on 117 genes detected using the summarized 5hmC profiles in cfDNA accurately distinguished HCC patients from controls (AUC = 95.1%; 95% CI, 93.6-96.5%) in the validation set, markedly outperformed α-fetoprotein (AFP) with superior sensitivity. The wd-scores, which not only detected early BCLC stages (e.g., Stage 0: AUC = 96.2%; 95% CI,94.1-98.4%) and small tumors (e.g., < 2 cm: AUC = 95.7%; 95% CI: 93.6-97.7%), also showed high capacity for distinguishing HCC from non-cancer patients with CHB/LC (AUC = 80.2%; 95% CI, 75.8-84.6%). Moreover, the prognostic value of 5hmC markers in cfDNA was evaluated for HCC recurrence, showing that a weighted prognostic score (wp-score) based on 16 marker genes predicted the recurrence risk (HR = 6.67; 95% CI, 2.81-15.82, p < 0.0001) in 555 patients who have been followed up after surgery. In conclusion, we have developed and validated a robust 5hmC-based diagnostic model that can be applied routinely with clinically feasible amount of cfDNA (e.g., from ~2-5 mL of plasma). Applying this new approach in the clinic could significantly improve the clinical outcomes of HCC patients, for example by early detection of those patients with surgically resectable tumors or as a convenient disease surveillance tool for recurrence.

## Introduction

Despite different therapeutic modalities currently available, patients with hepatocellular carcinoma (HCC) have undesirable outcomes, with five-year overall survival rates of less than 50% but can reach up to 70% with surgical resection or transplantation for early stage patients, highlighting the tremendous clinical need for effective early diagnostic approaches.(1, 2) HCC represents the second most common cause of cancer deaths worldwide (~750,000 annual deaths).(3) HCC is especially significant in East Asia, where major risk factors (e.g., chronic hepatitis B virus [CHB] infection and liver cirrhosis) are endemic. Indeed, China accounts for more than 50% of new HCC cases and related mortality.(4) Almost half of HCC patients have been diagnosed at an advanced stage, which often prevents the possibility of curative therapies. The early diagnosis of HCC can be difficult because of the absence of pathognomonic symptoms and the lack of sensitive and specific biomarkers, which is a main contributor to the poor outcomes of HCC patients.

At present, α-fetoprotein (AFP) is the most common serological test used for screening and diagnosis of HCC, as well as surveillance after treatment. However, there are serious limitations with serum-AFP-based diagnosis, such as low sensitivity,(5) possible false-negatives, and false-positives owing to confounding conditions.(6, 7) In addition, the unequivocal diagnosis of a nodule detected using ultrasonography remains clinically challenge and suffers unsatisfactory diagnostic accuracy.(8)

There are several challenges intrinsic to tissue pathology-based approaches such as high cost, limited tumor accessibility, and the issue of intra-tumoral heterogeneity.(9, 10) There are now some appealing alternatives, including methods based on liquid biopsy. Early examples of such methods targeted proteins or microRNAs, but these approaches have not typically offered satisfactory performance in terms of specificity and sensitivity or have not been suitable for deployment at the clinical scale.(11, 12) More recently, circulating cell-free DNA (cfDNA) in plasma has been associated with malignancy, systemic inflammation, and trauma.(13) Circulating cfDNA is known to carry genetic and epigenetic information from cells of origin.(14) Though limited by sample size and/or technical restrictions, several recent studies have begun to show the promise of epigenetic markers in cfDNA for diagnosis and prognosis in human cancers including HCC.(15, 16)

In the human genome, 5-hydroxymethylcytosines (5hmC) are abundant epigenetic marks that are generated through oxidation of 5-methylcytosines (5mC) by the ten-eleven translocation (TET) enzymes.(17) The 5hmC modifications occurring in promoters, regulatory elements, and in particular gene bodies faithfully reflect gene expression activation, thus being ideal markers for the activation of specific genes or genomic loci.(17) In this large clinical study, we employed our 5hmC-Seal, a highly sensitive and selective chemical labeling-based sequencing technology,(18, 19) to characterize 5hmC profiles in cfDNA from a cohort of ~3,000 subjects including HCC patients, controls and patients with other related diseases. We developed a weighted diagnostic score (wd-score) based on signature genes detected using the 5hmC profiles in cfDNA. In addition, the prognostic value of 5hmC markers in cfDNA was evaluated for predicting HCC recurrence risk, showing the potential of using 5hmC-Seal in cfDNA to develop prognostic markers.

## Methods

### Study Cohort

Informed consent was obtained from all participants prior to data collection, following the guidelines of the ethics committees of participating institutions. This study examined 2,728 individuals, including patients with HCC (n =1,208), controls (n = 965) (572 healthy individuals and 393 patients with benign lesions), as well as patients with chronic hepatitis B infection (CHB) (n = 291), liver cirrhosis (LC) (n = 110), and cholangiocarcinoma (CCC) (n = 154) (Figure 1, Supplemental Table 1), and met *a priori* defined quality criteria (i.e., at least 70% of annotated gene bodies with at least 30 read counts). Our primary training set was comprised of 1,718 randomly selected samples (806 HCC patients, 644 controls, and 268 CHB/LC patients). The remaining 402 HCC patients, 321 controls, and 133 CHB/LC patients comprised the validation set. The study subjects were recruited from Zhongshan Hospital, Fudan University, Shanghai, China; the Eastern Hepatobiliary Surgery Hospital, Shanghai, China; and Shanghai Public Health Clinic Center, China, from July 2016, to November 2017. In general, 5-10 mL of peripheral blood was collected for each individual, followed by plasma preparation and cfDNA extraction.(19) Detailed inclusion and exclusion criteria are provided in the Online Supplemental Content.

**Figure 1.**
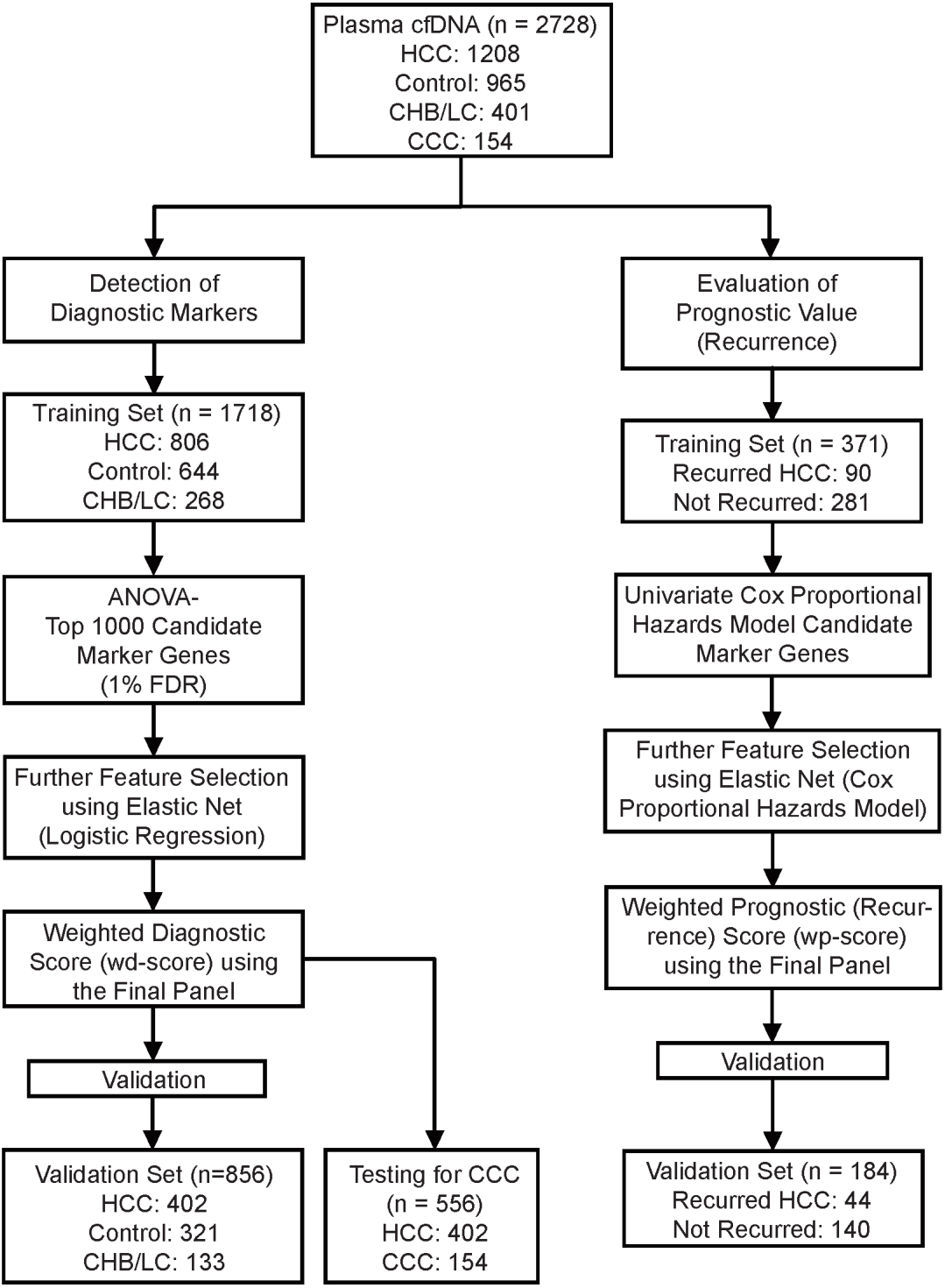
Study Design. The primary aim is to develop a weighted diagnostic score (wd-score) using the 5hmC-Seal data in plasma cfDNA that can distinguish patients with hepatocellular carcinoma (HCC) from controls (healthy individuals and patients with benign liver lesions), and from patients with chronic hepatitis B virus infection history (CHB)/liver cirrhosis (LC). The prognostic value of 5hmC markers in cfDNA is evaluated for predicting recurrence risk within 18 months after surgical resection. CCC: cholangiocarcinoma.

### Sample Preparation, 5hmC-Seal Profiling, and Data Processing

Detailed information on preparation of cfDNA from blood or genomic DNA (gDNA) from tissues, 5hmC-Seal library construction, sequencing, and data processing has been reported previously, (18, 19) and introduced in the Online Supplemental Content. Briefly, the 5hmC-Seal profiling combines a series of routine sample preparation and sequencing steps with a unique pull-down step based on covalent chemistry (Supplemental Figure 1) and is technically robust (Supplemental Figure2). The raw sequencing reads were then removed for adaptor sequences and checked for quality, followed by alignment to hg19.(19) High quality alignments were then counted for overlapping with gene bodies or other genomic features.

### Development of a Diagnostic Score for HCC

Detailed description of statistical analysis is shown in the Online Supplemental Content and Supplemental Figure 1. Briefly, in the top 1,000 candidate genes, we applied the elastic net regularization on a logistic linear regression model implemented in the *glmnet* package in the R language,(20) before the final multivariate logistic model was constructed. A weighted diagnosis score (wd-score) was then computed by multiplying the coefficients from above logistic model and the corresponding 5hmC value of marker genes for each participant:

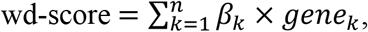

where *β_k_* is the coefficient from the final multivariate logistic model for the *k*th marker gene, and where *gene_k_* is the 5hmC value of *k*th marker gene. The AUC and 95% confidence intervals (CI) were generated to evaluate the performance of the wd-score in both training and validation sets. Sensitivity and specificity were estimated using the score cutoff that maximized the Youden’s index (*i.e*., sum of sensitivity and specificity).

### Exploration of a Prognostic Score for HCC Recurrence

We explored the prognostic value of 5hmC in cfDNA for HCC recurrence risk in 555 patients who underwent surgical resection and have been followed up within 18 months, using similar approaches as the wd-score computation (Supplemental Figure 1) based on the elastic net Cox model (Figure 1). A weighted prognostic (recurrence) score (wp-score) was then computed for each individual.

## Results

### Characterization of the 5hmC-Seal Profiles in cfDNA

We compared the 5hmC-Seal count data between duplicate samples from the same individual across different ranges of input DNA (i.e., 1 ng, 2 ng, 5 ng, and 10 ng) and two batches (Supplemental Figure 2). The 5hmC-Seal data showed high correlation (mean Pearson’s correlation > 0.992) between duplicate samples and different batches, demonstrating the robustness of this approach in clinically feasible amount of DNA materials (e.g., 2-10 ng from 2-5 mL of plasma). Overall, we did not observe obvious subpopulations within each group of cfDNA samples, indicating no systematic bias in the 5hmC-Seal profiling of these samples.

Comparing 26 pairs of tissue and plasma samples from the same individuals, the 5hmC profiles in cfDNA were shown to reflect tissue information (e.g., highly informative genes), thus supporting its use as a surrogate for tissue biopsy (Supplemental Figure 3). In a random set of cfDNA samples from 50 HCC patients and 50 healthy individuals, the distribution of 5hmC profiles was found to be enriched in gene bodies and depleted in the flanking regions (Supplemental Figure 4), consistent with previous findings.(19) Specifically, the detected 5hmC profiles in cfDNA samples from HCC patients were significantly enriched with liver tissue-derived histone modification marks for active gene expression, including H3K4me1 using the Roadmap Epigenomics Project data,(21) compared to healthy individuals (Supplemental Figure 4).

### Development of a Weighted Diagnostic Score for HCC

Our primary aim was to develop a convenient and integrated diagnostic score using 5hmC profiles in cfDNA to distinguish HCC patients from controls as well as from patients with CHB/LC. The multinomial elastic net approach suggested a consistent panel of 117 marker genes of 5hmC profiles (Supplemental Figure 1, Supplemental Table 2, Figure 2A-B), the wd-scores (range: 23.51-26.31; mean = 24.95, sd = 0.33) computed based on which showed excellent capacity for distinguishing HCC patients from controls in both training set (AUC = 97.2%; 95% CI, 96.5-97.9%) and validation set (AUC = 95.1%; 95% CI: 93.6-96.5%) (Figure 2C-D), or from benign lesions only (Supplemental Figure 5), markedly outperformed AFP (Training Set: AUC = 85.5%; 95% CI: 83.5-87.5%; Validation Set: AUC = 84.7%; 95% CI: 81.8-87.6%) (Figure 2C-D). Specifically, at the cutoff of 24.93, the wd-scores achieved the highest Youden’s index with a sensitivity of 88.7% and a specificity of 93.2% in the training set, and a sensitivity of 83.8% and a specificity of 91.3% in the validation set (Figure 2C-D). While AFP showed a high specificity at both 20 ng/mL and 400 ng/mL in distinguishing HCC patients from controls, our wd-scores demonstrated an overall higher diagnostic performance, for those HCC patients misclassified by AFP (e.g., at 20 ng/mL, 339 HCC patients were misclassified by AFP in the training set) (Supplemental Figure 6).

**Figure 2.**
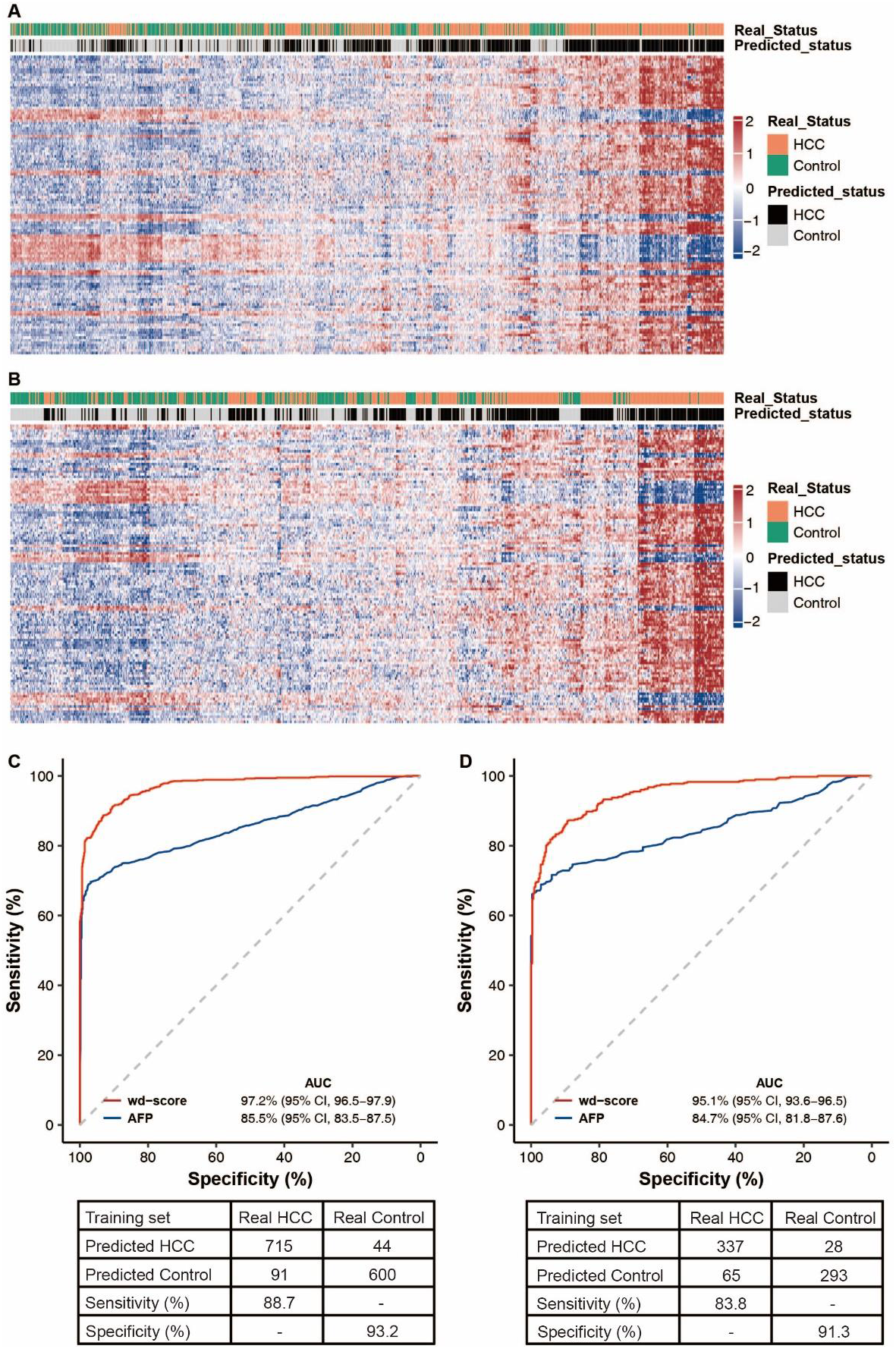
Performance of the 5hmC Diagnostic Model for HCC. The 117 genes used to compute the wd-scores for HCC diagnosis are used to generate the heatmaps in (A) training and (B) validation set, respectively. Predictive performance of the wd-scores in distinguishing HCC from controls is shown for (C) training and (D) validation set, respectively. At the cutoff of 24.93, the wd-scores achieved the highest Youden’s index.

Individuals with a history of CHB infection are at 5-100 fold higher risk for developing HCC.(22) Consistent with HCC epidemiology in China, 85.0% of the HCC patients in our study cohort had a history of CHB infection. Liver cirrhosis is one major risk factor for HCC, and cirrhosis frequently complicates HCC diagnoses. In our study, 65.9% of the HCC patients had pathologically confirmed cirrhosis. Notably, the 117 gene-based wd-scores also distinguished HCC patients from patients with CHB/LC in both training set (AUC = 80.7%; 95% CI: 77.7-83.6%) and validation set (AUC = 80.2%; 95% CI, 75.8-84.6%) (Supplemental Figure 7). Overall, the wd-scores showed no strong correlation with AFP across different sample groups and were significantly higher in HCC patients than in CHB/LC patients or controls (t-test p<0.001) (Figure 3A-B), indicating independent predictive capability of this integrated diagnostic algorithm. Finally, CCC, which has different cell origin, risk factors, and prognosis, is also prevalent, representing ~10-15% of all diagnosed primary liver cancers. Interestingly, though not trained with CCC, our wd-scores showed a strong trend of separation (AUC = 78.0%; 95% CI: 73.9-82.2%) between the 402 HCC patients in the validation set and the 154 CCC patients recruited in our study cohort (Supplemental Figure 8).

**Figure 3.**
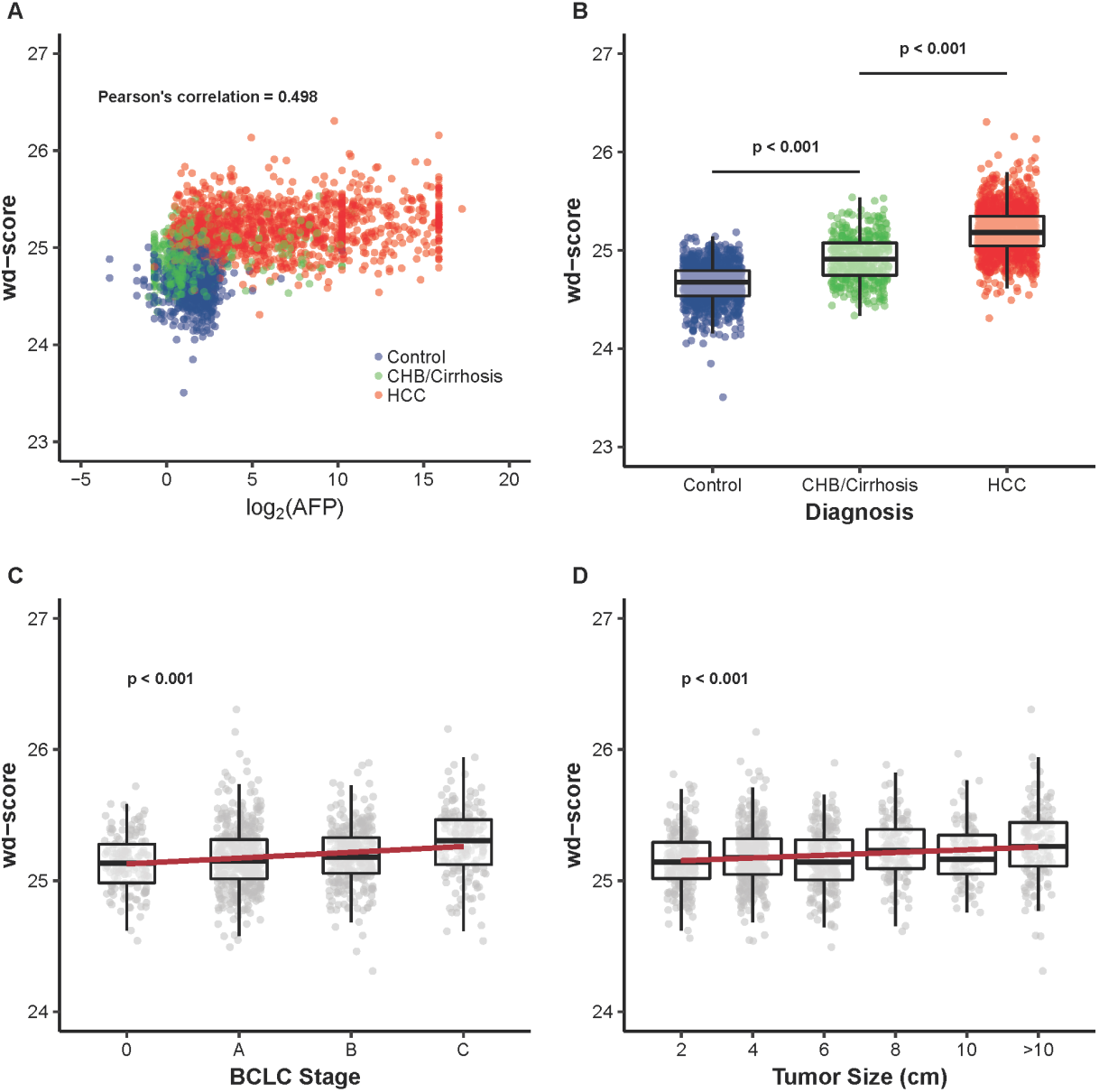
Relationships between the wd-scores and Clinical Characteristics. The wd-scores computed using the 117 marker genes for HCC diagnosis are investigated for their relationships with clinical characteristics, including (A) AFP level; (B) diagnosis; (C) BCLC stage; and (D) tumor size.

Further functional annotation analysis of the 117 marker genes suggested an enrichment of pathways “glycolysis/gluneogenesis” (*G6PC*, *ALDOB*, *ADH1B*, *ALDH3A2*, and *PCK1*) and “retinol metabolism” (*CYP26C1*, *CYP2C18*, *HSD17B6*, and *ADH1B*) (Supplemental Figure 9A), relevant to various liver functions and liver impairment in patients with CHB/LC and liver cancer.(23, 24) Notably, these 117 genes were enriched with genes specifically expressed in liver tissue (hypergeometric p<0.0001) (Supplemental Figure 9B), indicating their tissue relevance. Moreover, hierarchical clustering showed that the distance between the differential profiles of liver plasma and liver tissue was shorter than those derived from colon and stomach cancer patients,(19) suggesting cancer-type specificity of the differential 5hmC profiles (Supplemental Figure 10).

### Diagnostic Scores and Clinical Characteristics

We then examined the 117-gene wd-scores for individual HCC patients by BCLC (Barcelona Clinic Liver Cancer) stages (Figure 3C). We found that the wd-scores increased as BCLC stage advanced in all HCC patients with available stage information (n = 1,036) (effect size = 0.255, p-trend < 0.001). Specifically, late stages (Stage B/C) showed higher wd-scores than early stage patients (Stage 0/A) (t-test p<0.001). The predictive performance of wd-scores in the training and validation sets by the BCLC stage is shown in Supplemental Figure 11. Notably, the wd-scores not only detected late stage HCC patients, but also showed excellent predictive power for early stage patients, specifically for Stage 0 patients (AUC = 96.2%; 95% CI: 94.1-98.4%) and Stage 0/A patients (AUC = 94.1%; 95% CI: 92.1-96.0%) in the validation set.

Further, while we are fully aware that tumor size is not a highly meaningful indication of progression in HCC, we examined the correlation between measured tumor sizes and the wd-scores and noted a strong trend of association in all HCC patients with confirmed tumor size information (n =1,113) (effect size = 0.109; p-trend < 0.001) (Figure 3D). Note that this 117-gene wd-score accurately distinguished patients with different tumor size ranges (from < 2 cm to > 10 cm) from controls in both training set and validation set (Supplemental Figure 12). Specifically, for HCC smaller than 2 cm (n = 224), the wd-scores showed a strong predictive performance by separating HCC from controls in the validation set (AUC = 95.7%; 95% CI: 93.6-97.7%). In addition, the wd-scores showed equivalent performance of distinguishing CHB/LC (or benign lesions only) from either small HCC (< 2 cm) or early stage patients (Stage 0/A) in the validation set, as compared to the accuracy of separating between all HCC and CHB/LC (Supplemental Figure 5, 7).

### Evaluation of a Prognostic Score for Recurrence Risk

We then analyzed 5hmC data in cfDNA from 555 HCC patients who have been followed up after surgery at our participating hospitals (Supplemental Table 3). In total, 134 HCC patients recurred and 421 patients did not recur within 18 months. Specifically, a 16-gene signature for recurrence risk (Supplemental Figure 13, Supplemental Table 4) was detected in the training set (Recurred: n = 90; Not recurred: n = 281; Figure 1). We then explored the prognostic value of the wp-score computed using this 16-gene recurrence signature in HCC patients (Figure 4). The HCC patients were successfully grouped by differential recurrence risk based on the median of the wp-scores in both training set (HR = 5.73; 95% CI: 3.41-9.64; log-rank p < 0.0001) and validation set (HR = 6.67; 95% CI: 2.81-15.82; log-rank p < 0.0001). Moreover, the wp-scores for recurrence in cfDNA showed a predictive performance that complemented BCLC stage alone (log-rank p < 0.0001) (Supplemental Figure 14). Interestingly, combing stage information and the 16-gene wp-score, a separate subgroup of HCC patients with a distinct recurrence risk was detected. Specifically, Stage 0/A patients with a high wp-score as well as Stage B/C patients with a low wp-score displayed an intermediate risk for recurrence (Figure 4C-D). For example, this newly detected subgroup with intermediate risk showed an HR of 6.29 relative to the low risk group, significantly different from the high risk group (HR = 33.73) in the validation set. Notably, exploring these 16 genes for recurrence risk using the Human Pathology Atlas (HPA)(25) for liver cancer five-year survival showed that four of them were previously associated with five-year survival in the HPA (log-rank p < 0.05), including *CCNY*, *TALDO1*, *TMEM183A*, and *UQCRH*. Besides the HPA, many recurrence-associated genes have been implicated in cancer survival, progression, or metastasis, e.g., *PAX8*, *YY1AP1*, *CAPN2*, and *LIMK2*.(26-29)

**Figure 4.**
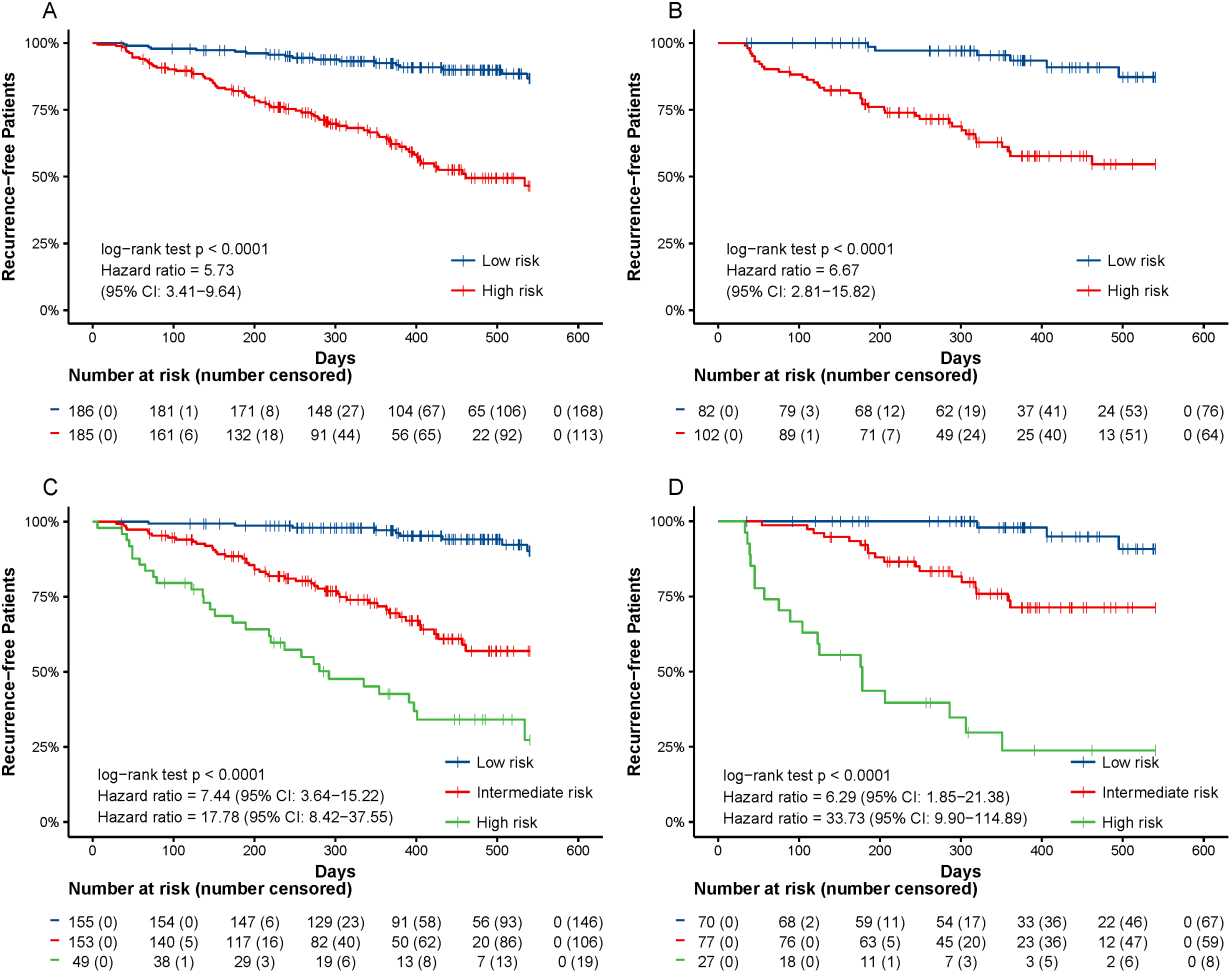
Prognostic Value of 5hmC for HCC Recurrence. The Kaplan-Meier (KM) analysis was performed for HCC patients who have been followed up within 18 months after surgery, based on the wp-scores computed using the 16 marker genes for recurrence risk in (A) training and (B) validation set, respectively. The wp-scores show complementary prognostic value with BCLC stages by identifying a subgroup of patients with a unique recurrence risk (intermediate) in (C) training and (D) validation set, respectively.

## Discussion

We sought to identify and validate clinically convenient, liquid biopsy-based epigenetic biomarkers to help diagnose HCC patients from controls and related liver diseases using the 5hmC-Seal, a highly sensitive chemical labeling technique in cfDNA. Our primary analysis identified a 117-gene based 5hmC marker panel and algorithm (wd-score) that not only displayed significantly improved performance than diagnosis (HCC vs. controls) based on serum AFP levels (Figure 2), but also showed excellent capacity for distinguishing HCC from CHB/LC, the two major related liver diseases that often complicate HCC diagnosis. Notably, the 117-gene based wd-score successfully detected HCC patients in those patients failed to be detected by different AFP cutoffs (Supplemental Figure 6), suggesting superior sensitivity of the wd-score over AFP. Our results further established the diagnostic performance of the 117-gene based wd-score in cfDNA by distinguishing HCC patients at different BCLC stages from controls (Supplemental Figure 11) or CHB/LC (Supplemental Figure 7), including notably patients at early stage HCC (Stage 0/A) from other liver diseases. For HCC patients with different tumor sizes, the wd-score demonstrated consistently high capacity for distinguishing HCC patients from controls (Supplemental Figure 12). Taken together, our findings suggested that as a robust, non-invasive epigenetics-based approach, the 5hmC-Seal could become an integrated part of HCC management, from detection of early stage patients or patients with small tumors (e.g., < 2 cm)through real-time post-treatment surveillance. We envision that patients diagnosed positive with 5hmC markers can be put on more frequent monitoring by ultrasonography and CT/MRI, thus improving patient survival in the long run. Given the whole-genome nature of the 5hmC-Seal assay, the convenience of the “one-for-all” 117-gene based wd-score could be complemented by various pair-wise models developed to distinguish specifically, for example HCC and CHB/LC (Supplemental Figure 15; Supplemental Table 5), thus providing an alternative approach to improve diagnosis in specific clinical situations at higher precision.

Clinically, for those patients showing a high risk of recurrence, they could receive adjuvant transarterial chemoembolization and adjuvant molecule-targeted drugs (e.g. sorafenib), and be subject to more frequent follow-up. We, therefore explored their potential prognostic value for predicting recurrence risk using the 5hmC-Seal data obtained in cfDNA before surgical resection (Figure 4A-B; Supplemental Figure 13-14). We showed that the 5hmC-Seal data form cfDNA could be used to complement conventional prognostic tools, such as the BCLC staging system. Our recurrence-associated 5hmC markers in cfDNA could further stratify these patients by identifying those with a unique recurrence risk profile that otherwise could not be explained by stage alone (e.g., lower risk among the advanced patients), thus providing opportunities in the future for more precise care of this distinct patient subgroup (Figure 4; Supplemental Figure 14). Note that the follow-up period in the current study was relatively short (18 months), a long-term follow-up study will be performed to further clarify the prognostic value of 5hmC markers, though an evaluation using the recent HPA data(25) on liver cancer five-year survival did indicate prognostic relevance of our recurrence-associated 5hmC markers.

Technically, in contrast to other screening technologies, the 5hmC-Seal approach does not require pathology-related assumptions to inform probe targeting strategies. Because of its covalent chemical labeling nature which prevents sequencing biases, the 5hmC-Seal approach is not limited to any specific sequence context.(18) Further, given that cfDNA is highly fragmented (~160-320 bp), our approach of summarizing 5hmC profiles in cfDNA using genomic “bins” (*i.e*., gene bodies) is robust in terms of noise and signal as well as convenient for functional exploration.(19) The 5hmC-Seal data are not limited to specific sites, avoiding the potential problem of missing specific sites encountered in mutation-based analysis.(30)

Regarding public health importance, notable utilities of the 5hmC markers in cfDNA for HCC include: i) even though screening of patients with established cirrhosis with ultrasonography every 6 months is recommended for early diagnosis and improved overall survival, ultrasonography has only a sensitivity of 60-80%.(31) Our sensitive 5hmC markers in cfDNA for HCC (Figure 3; Supplemental Figure 11-12) could fill an urgent unmet need to identify small tumors and/or early stage tumors amenable to curative treatments in order to improve survival. Thus we offered a significantly improved tool for early detection of HCC, especially in patients with cirrhosis, for which a confident diagnosis of nodules is almost impossible;(32) and ii) given its flexibility, robustness, and non-invasiveness, the 5hmC-Seal approach may also be used for public health screening of high-risk populations or even the population at large. Particularly, current diagnostic imaging techniques for HCC require a combination of equipment and infrastructural support which might not be readily available in developing regions where HCC burden is extremely high.

We acknowledge several limitations. Firstly, we used a case-control design in the current study, a clinical trial for predictive marker validation will be required in the future to further evaluate the diagnostic power of our model. Secondly, although the major clinical variables (e.g., gender, age group) were well-balanced between the validation sets and the original training sets (Table A1), the validation sets could have some selection biases. Thirdly, our study was in a Chinese patient cohort mostly with CHB/LC background, and therefore more validation is warranted to demonstrate the generalizability of the results in prospective studies which will cover other populations, geographical regions, and disease risk factors.

In conclusion, we have developed and validated a robust 117-gene diagnostic model for HCC using a sensitive assay (5hmC-Seal) in cfDNA. Applying this new approach in the clinic could significantly improve the clinical outcomes of HCC patients by identifying those patients with surgically resectable tumors or those with distinct clinical subgroups.

